# Are reads required? High-precision variant calling from bacterial genome assemblies

**DOI:** 10.1101/2025.02.20.639404

**Authors:** Ryan R. Wick, Louise M. Judd, Timothy P. Stinear, Ian R. Monk

## Abstract

1.

Accurate nucleotide variant calling is essential in microbial genomics, particularly for outbreak tracking and phylogenetics. This study evaluates variant calls derived from genome assemblies compared to traditional read-based variant-calling methods, using seven closely related *Staphylococcus aureus* isolates sequenced on Illumina and Oxford Nanopore Technologies platforms. By benchmarking multiple assembly and variant-calling pipelines against a ground truth dataset, we found that read-based methods consistently achieved high accuracy. Assembly-based approaches performed well in some cases but were highly dependent on assembly quality, as errors in the assembly led to false-positive variant calls. These findings underscore the need for improved assembly techniques before the potential benefits of assembly-based variant calling – such as reduced computational requirements and simpler data management – can be realised.

**Impact statement:** Variant calling is foundational to microbial genomics, yet traditional workflows rely heavily on sequencing reads, which for a typical bacterial genome can be hundreds of megabytes. In contrast, genome assemblies are far smaller – usually just a few megabytes – making them significantly easier to manage. If accurate variant calls could be made directly from assemblies, this would reduce computational demands and, in some cases, may even eliminate the need to retain raw sequencing reads. This study addresses the key question of whether variant calling from assemblies is accurate enough to replace read-based methods. Our findings show that while assembly-based variant calling can achieve high accuracy, this is only possible with error-free assemblies. Since most assemblies contain errors, assembly-based variant-calling approaches should currently be used with caution. Nevertheless, as sequencing and assembly technologies continue to advance, improved assembly accuracy may make assembly-based variant calling a viable alternative, reducing data complexity and storage demands while streamlining microbial genomic analyses.

**Data summary:** Supplementary methods, data, figures and tables are available at github.com/rrwick/Are-reads-required which is also archived on Zenodo (10.5281/zenodo.14868870). Reads and assemblies are available via BioProject PRJNA1193226.

## 4. Introduction

Accurate identification of genetic variants is crucial in microbial genomics, particularly in applications such as building phylogenies, tracking outbreaks and studying evolutionary processes.^1,2^ Single nucleotide polymorphisms (SNPs) are often the primary focus of variant-calling efforts, but other variants, such as insertions and deletions (indels), can also provide valuable information.^3^ High-precision variant calls (i.e. with few to no false positives) are especially important for closely related samples, where even small differences, such as 10 SNPs, can determine whether or not bacterial isolates are considered part of the same outbreak.^4^

In many ways, genome assemblies offer significant advantages over whole-genome sequencing reads: they have smaller file sizes, are easier to manage and are conceptually simpler. Long-read sequencing, such as from Oxford Nanopore Technologies (ONT) platforms, can usually enable complete genome assemblies, where each genomic element is assembled into a single contiguous sequence, revealing the large-scale structure of the genome.^5^ Given these benefits, it would be convenient to extract variants directly from assemblies, removing the need to store and process reads. However, the question remains: can assembly-based variant-calling achieve the level of accuracy required for microbial analyses?

To explore this question, we investigated the accuracy of variant calls from both read-based and assembly-based methods from a series of *Staphylococcus aureus* suppressor mutants. These isolates were sequenced using both Illumina short reads and ONT long reads. By comparing multiple variant-calling approaches, we aimed to determine whether read-based methods are necessary for precise microbial variant detection, or whether assembly-based approaches can provide similarly accurate results.

## 5. Methods

In this study, we used seven closely related bacterial isolates. The parental strain NRS384 is a USA300 community-associated methicillin-resistant clone of *S. aureus*. A targeted mutant was constructed in the essential histidine kinase WalK (yielding WalK^T389A^) to inactivate its phosphatase activity and prevent dephosphorylation of the essential cognate response regulator WalR.^6^ Propagation of a single colony of WalK^T389A^ aerobically (200 rpm) at 37°C in 10 ml of Brain Heart Infusion broth (BHI, Oxoid) yielded suppressor mutants at a high frequency which could be distinguished from WalK^T389A^ by a change in haemolytic activity on Sheep Blood agar (5% Sheep blood in Columbia agar, SBA), Atl activity (BHI agar (BHIA) containing 0.2% *Micrococcus luteus* cells (Sigma, M3770), MLA) and/or pigmentation on Mueller-Hinton agar (MHA, Difco) (**Figure S1**). Suppressors (sample names beginning with IMAL) were single-colony purified on BHIA and stored at -80°C. Genomic DNA isolation and phenotypic analysis on SBA and MLA were conducted as described previously.^6^

ONT sequencing was performed on an R10.4.1 MinION flow cell using the native barcoding kit (SQK-NBD114-96) and then basecalled using Dorado v0.7.3 using the sup@v5.0.0 model. ONT reads were quality controlled by discarding any read with an average qscore (as reported by Dorado) of less than 12.5 or a length of less than 1 kbp. Illumina sequencing was performed on an Illumina NextSeq 2000 with a 150 bp PE kit. Illumina reads were quality controlled using fastp^7^ v0.23.4 with default parameters. See **Table S1** for read details and accessions.

For each of the seven genomes, we generated carefully curated ground-truth assemblies using Trycycler^5^ v0.5.5, Polypolish v0.6.0 and Pypolca^8^ v0.3.1, and each assembly was reorientated to a consistent starting position using Dnaapler^9^ v0.8.1. To check for potential errors, we ran Clair3^10^ v1.0.10 (using ONT reads), Sniffles2^11^ v2.4 (using ONT reads) and Freebayes^12^ v1.3.8 (using Illumina reads) to screen for variants (i.e. potential errors) in the assemblies, each of which found no discrepancies. Each assembly contained a 2.88 Mbp chromosome (which differed slightly across the seven genomes) and two small plasmids (4439 bp and 3125 bp, which were identical across all seven genomes). Trycycler v0.5.5 was used to generate a multiple sequence alignment of all seven chromosomes, and invariant sites were then removed to produce the alignment shown in **Figure 1**.

**Figure 1:**
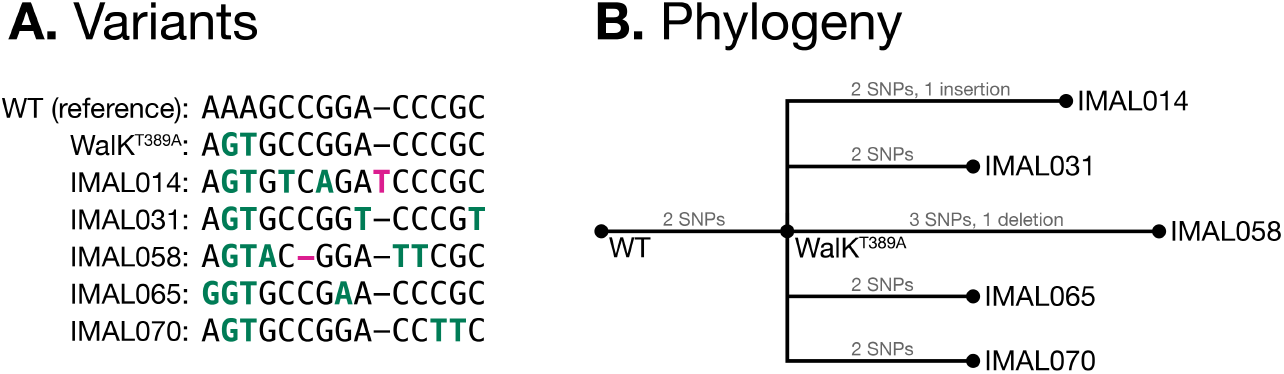
The relationship between the seven *S. aureus* NRS384 isolates used in this study. **A:** an invariant-free multiple sequence alignment, showing all variants between the isolates and the wild-type (WT) reference: 23 SNPs (green) and 2 indels (pink). **B:** the phylogenetic relationship between the isolates, as constructed by IQ-TREE v2.3.6^1^.

For each isolate, we used Rasusa^13^ v2.1.0 to randomly subsample both the ONT and Illumina read sets to 20× (low), 50× (medium) and 100× (high) depths. Each subsampled read set was then used to call variants against the reference assembly (wild-type NRS384), using Clair3 v1.0.10 for ONT reads and Freebayes v1.3.8 for Illumina reads. For Clair3, variants were kept that had filter status of ‘PASS’.^14^ For Freebayes, variants were kept that had a quality score ≥100 (the threshold used by Snippy^15^). See **Supplementary methods** for the exact commands used.

Each subsampled read set was then assembled using a variety of methods. Shovill v1.1.0, Unicycler^16^ v0.5.1 and SKESA^17^ v2.5.1 were used for short-read-only assemblies (both Shovill and Unicycler serving as wrappers for SPAdes^18^ v4.0.0). Canu^19^ v2.2, Flye^20^ v2.9.5 and Raven^21^ v1.8.3 were used for long-read-only assemblies. Unicycler v0.5.1 and Hybracter^22^ v0.9.0 were used for hybrid (both short- and long-read) assemblies. Long-read-only assemblies were additionally polished using Medaka^23^ v2.0.0 using its bacterial model.

Variants were called from each assembly (all assemblies of subsampled reads as well as our ground-truth assemblies) using three different methods (**Figure 2**). The first method, which we abbreviate as ‘MUMmer’, uses the dnadiff tool from MUMmer^24^ v4.0.0rc1 to directly align the assembly to the reference genome and identify differences, which were then converted to VCF format using all2vcf^25^ v0.7.8. The second method, we which abbreviate as ‘Shred’, uses wgsim to generate synthetic reads (150 bp, error-free, 100× depth), aligns them to the reference with BWA MEM^26^ v0.7.18 and then calls variants using Freebayes v1.3.8 (with the same ≥100 quality threshold used in our read-based variant calls). The third method uses the ska map command from SKA2^27^ v0.3.11 to find variants against the reference. Each of these methods is bundled into an easy-to-run script available in **Supplementary methods**.

**Figure 2:**
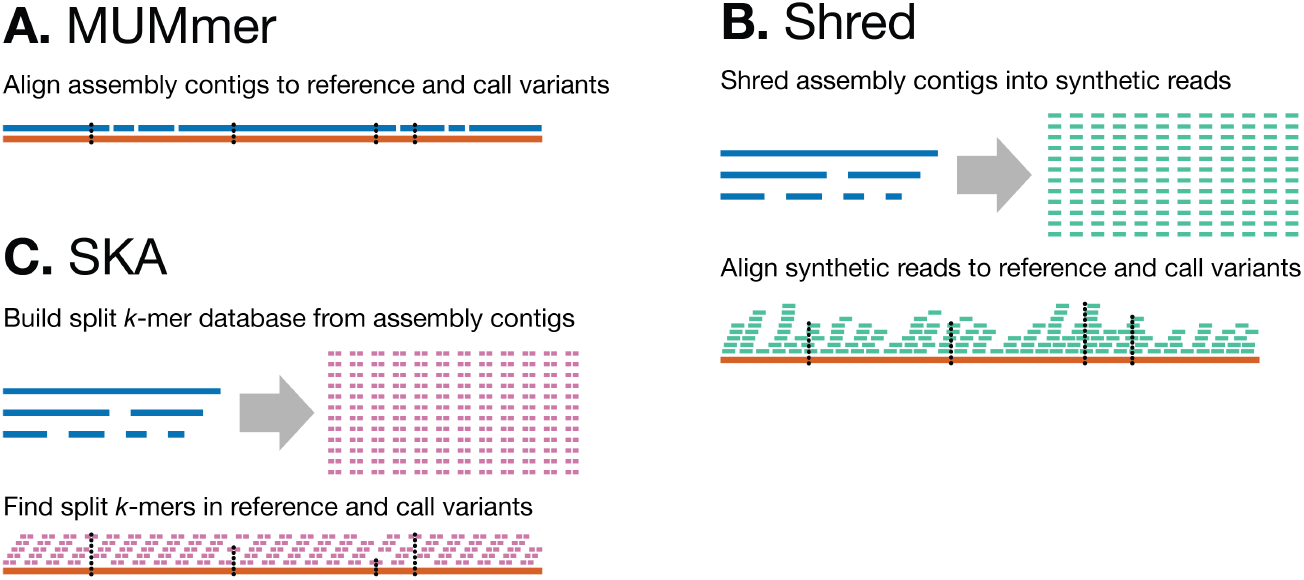
Methods for calling variants from assemblies. **A:** The MUMmer method relies on directly aligning the assembly’s contigs to the reference genome, with variants called from these alignments. **B:** The Shred method produces synthetic error-free reads from the assembly’s contigs and aligns them to the reference genome, allowing for the use of read-based variant callers such as Freebayes. **C:** The SKA method uses split *k*-mers to efficiently find SNPs between the assembly’s contigs and reference genome.

In total, this produced 756 VCF files: 42 from read-based variant calling (2 read types × 3 depths × 7 isolates), 21 from reference assemblies (3 variant-calling methods × 7 isolates), 189 from short-read assemblies (3 assembly methods × 3 depths × 3 variant-calling methods × 7 isolates), 378 from long-read assemblies (3 assembly methods × 3 depths × 3 variant-calling methods × 2 polishing methods [no polishing and Medaka] × 7 isolates) and 126 from hybrid assemblies (2 assembly methods × 3 depths × 3 variant-calling methods × 7 isolates).

To establish a truth set of variants, we built a whole-genome multiple sequence alignment using each of the variant-calling methods by applying the variants to the reference genome using BCFtools^28^ v1.21 and Trycycler v0.5.5. The following methods all produced the same results: Illumina-read Freebayes (all read depths), ONT-read Clair3 (100× depth), ground-truth assembly Shred and MUMmer, and Hybracter assembly Shred and MUMmer (all read depths). Since all of these methods produced the same variants, we considered these to be the ground-truth variants against which all methods were then assessed (one ground-truth VCF for each of the seven isolates). Totalled across all seven isolates, there were 23 ground-truth SNPs and 2 ground-truth indels (**Figure 1A**).

To quantify the accuracy of the variant calls, we used vcfdist^29^ v2.5.3 to compare each VCF file to its corresponding ground-truth VCF file. The following classification metrics were counted for both SNPs and indels and stored in **Table S2**: true positives (TP), false negatives (FN) and false positives (FP). This allowed for the calculation of sensitivity (*TP*/(*TP* +*FN*), the probability that a true variant will be called) and precision (*TP*/(*TP* + *FP*), the probability that a called variant is true).

## 6. Results

### 6.1 Assembly-based variant-calling method

We first assessed which of the three methods for assembly-based variant calling performed best, with **Table 1** showing SNP and indel metrics based on all 714 assembly-based VCF files. Both MUMmer and Shred had perfect sensitivity, i.e. they never failed to call an existing SNP or indel. SKA failed to call SNPs in close proximity to other variants and could not call indels, both limitations of its split-*k*-mer-based method. While MUMmer and Shred performed similarly, Shred produced slightly fewer false positives for both SNPs and indels. We therefore chose to only use Shred-based variant calls for our subsequent results, discarding MUMmer- and SKA-based variant calls for the subsequent analyses.

**Table 1:** True positive (TP), false negative (FN) and false positive (FP) variant calls for each of the assembly-based variant-calling methods, for both SNPs and indels, along with sensitivity (Sens) and precision (Prec).

|  | SNPs |  |  |  |  | Indels |  |  |  |  |
| --- | --- | --- | --- | --- | --- | --- | --- | --- | --- | --- |
|  | TP | FN | FP | Sens | Prec | TP | FN | FP | Sens | Prec |
| **MUMmer** | 782 | 0 | 4669 | 1.0 | 0.14 | 68 | 0 | 719 | 1.0 | 0.086 |
| **Shred** | 782 | 0 | 4001 | 1.0 | 0.16 | 68 | 0 | 699 | 1.0 | 0.089 |
| **SKA** | 374 | 408 | 7431 | 0.48 | 0.048 | 0 | 68 | 0 | 0.0 | n/a |

### 6.2 Read- and assembly-based variant calling

Using only the best-performing assembly-based variant-calling method (Shred), we then compared read-based and assembly-based variant calls (**Figure 3**). Overall, the best results came from Illumina-read variant calls (Freebayes) and Hybracter-assembly variant calls, both of which were error-free. ONT-read variant calls (Clair3) contained no false positives but did miss some SNPs at the 20× and 50× read depths (**Figure S2**). All other methods were prone to false-positive calls for both SNPs and indels.

**Figure 3:**
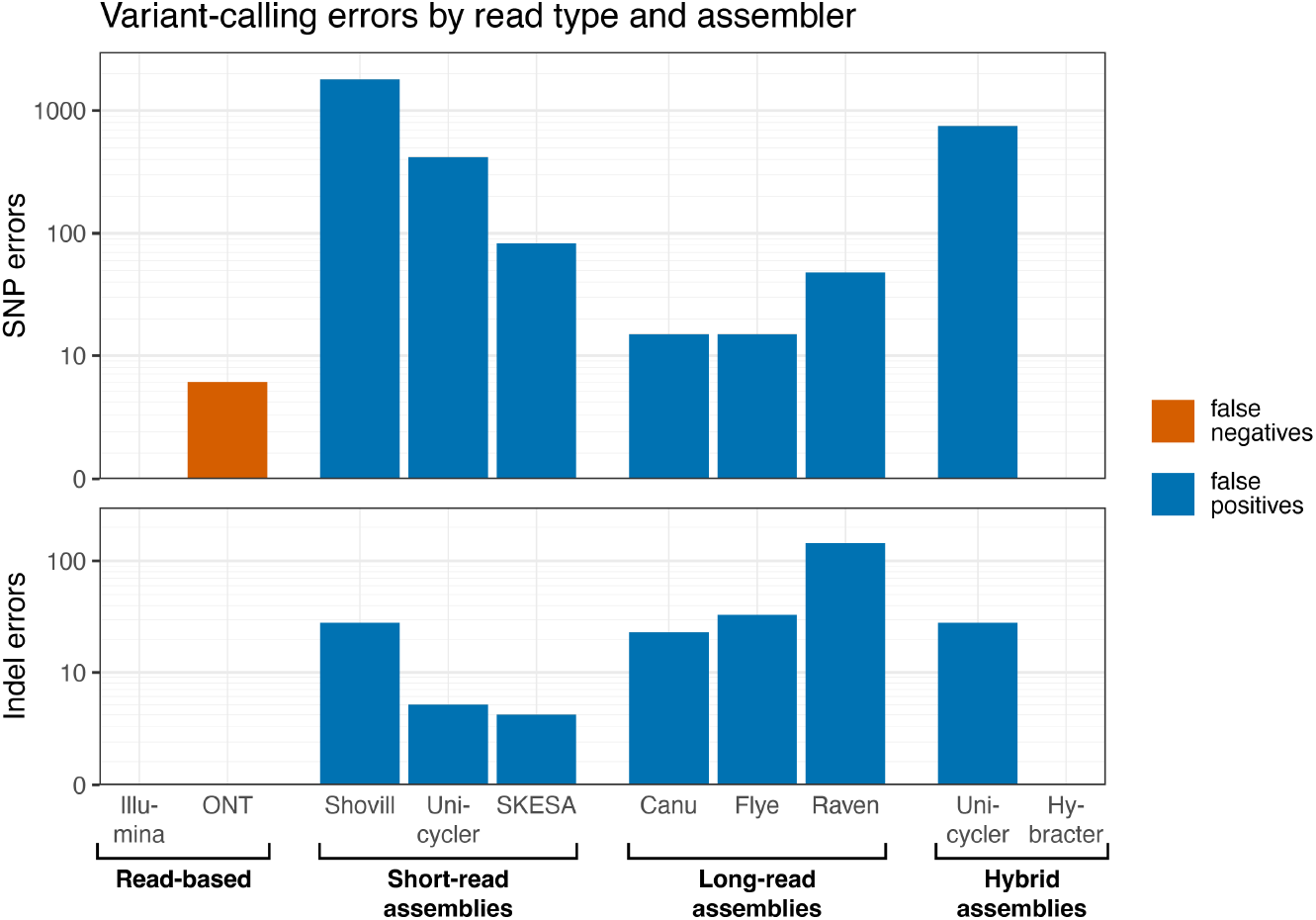
Variant-calling errors for both read-based and assembly-based variant-calling methods. For each method, the errors were summed across all genomes and read depths. See **Figure S2** for the results for each depth separately and for sensitivity and precision values. The y-axes have a pseudo-log transformation.

Short-read assemblies were particularly prone to SNP errors while long-read assemblies were more prone to indel errors. SKESA was the best-performing short-read assembler, likely due to its conservative heuristics (it prefers breaking contigs over assembling uncertain sequence). Canu was the best-performing long-read assembler, though Flye did nearly as well. For hybrid assemblers, there was a marked difference between Unicycler (which uses a short-read-first assembly method) and Hybracter (which uses a long-read-first assembly method), the latter performing much better for both SNPs and indels (**Figure 3**).

### 6.3 Medaka polishing

For each of the long-read assemblies, we performed polishing with Medaka in an attempt to improve the sequence accuracy. Surprisingly, this caused a large increase in the total number of variant-calling errors (**Figure S3**). Further investigation revealed these Medaka-introduced errors were primarily caused by two factors. The first was caused by an 850 bp region of homologous sequence shared between the *S. aureus* chromosome and the 4.4 kbp plasmid. The inability to assemble small plasmids is a common error in long-read assemblies,^30^ leading to the absence of the 4.4 kbp plasmid in some assemblies. When this occurred, Medaka’s alignment of reads to the assembly resulted in plasmid reads erroneously aligning to the chromosome, causing errors in the consensus algorithm.

The second factor related to increasing Medaka errors was when the assembly contained duplicated sequence from the genome. This occurred when the start and end of a circular contig overlapped and when a region of the genome was assembled into both a primary and alternative sequence. When these issues occur, Medaka was prone to introducing errors in the duplicated regions, leading to false-positive assembly-based variant calls. Canu is prone to both start-end overlaps and alternative-sequence contigs (what it calls ‘bubbles’), leading to the largest increase in variant-calling errors after Medaka polishing.

In assemblies where neither of these factors were present, we found Medaka often improved the accuracy of assembly-based variant calls (**Table S3**). This suggests that Medaka-polishing can be beneficial for variant calling when the assembly is correctly finished and contains the entire genome with no duplicated sequence.^31^

## 7. Discussion

In this study, we examined variant calling in a group of closely related *S. aureus* isolates which only had 25 combined variants across six mutant strains relative to the WT strain. We found that it was possible to obtain precise variant calls from genome assemblies (i.e. without directly using the reads), but this required very high-quality assemblies: only our ground-truth assemblies (Trycycler with manual curation) and Hybracter assemblies (a long-read-first hybrid assembly pipeline) yielded perfect variant calls. All other assembly methods resulted in false-positive calls, sometimes exceeding 100 errors per genome, leading to poor precision. This suggests that most genome assemblies are not suitable for variant calling if high precision is required.

In general, we found that short-read assemblies were more prone to false-positive SNP calls while ONT assemblies were more prone to false-positive indel calls. Our ONT assemblies were based on the latest ONT chemistry and basecalling model, and it is likely that older ONT assemblies will have reduced precision for both SNPs and indels. While we expected Medaka polishing to improve assembly-based variant calls for ONT assemblies, missing or duplicated sequences in the assemblies often led to Medaka introducing errors. This emphasises the need for an assembly to be structurally correct before Medaka polishing.

While we compared three different methods for assembly-based variant-calling (MUMmer, Shred and SKA), we did not attempt to optimise any of these methods. This is a key limitation of this study, and so higher precision assembly-based variant-calling methods are likely possible.

Such optimisations might include masking parts of the reference genome (e.g. repeats and low-complexity regions) or excluding parts of the assembly (e.g. short contigs and low-quality-bases^32^). If variant-calling from assemblies is necessary (e.g. because reads are not available), then further work is required to determine the optimal method. Since SKA’s split-*k*-mer approach cannot identify indels or closely spaced SNPs, the optimal assembly-based variant-calling method will likely involve alignment.

The *S. aureus* genomes used in this study were relatively easy to assemble – our Illumina assemblies often contained fewer than 100 contigs (**Table S2**) and error-free ONT assemblies were possible (**Table S3**). More challenging genomes (e.g. containing more repetitive regions) would likely make assembly-based variant calling even more challenging. Of the 30 samples assembled in the Hybracter manuscript,^22^ 12 had zero errors in their assembly while 18 contained errors. This suggests that even when performing a robust Hybracter hybrid assembly, false-positive assembly-based variant calls may occur for approximately half of bacterial genomes. Also, the *S. aureus* genomes in this study contained no structural differences relative to the reference genome. When structural differences occur, there is the possibility of more variant-calling errors for both read- and assembly-based approaches.

In conclusion, we found that assembly-based variant calling is prone to false-positive errors, so traditional read-based variant calling is preferred when possible. However, error-free assemblies can produce error-free variant calls, so as sequencing and assembly methods become more reliable in the future, assembly-based variant calls may become more feasible.

## 8. Author statements

### 8.1 Author contributions

Conceptualization: RRW, LMJ, IRM, Data curation: RRW, LMJ, Formal analysis: RRW, Funding acquisition: RRW, IRM, TPS, Investigation: RRW, LMJ, IRM, Methodology: RRW, LMJ, IRM, Project administration: IRM, TPS, Software: RRW, Resources: LMJ, IRM, Supervision: IRM, TPS, Visualization: RRW, Writing – original draft: RRW, Writing – review & editing: RRW, LMJ, TPS, IRM.

### 8.2 Conflicts of interest

The authors declare that there are no conflicts of interest.

### 8.3 Funding information

RRW is supported by an ARC Discovery Early Career Researcher Award (DE250100677). TPS is supported by NHMRC Research Fellowship (APP1105525) and ARC Discovery Project (DP240102465).

## 8.4 Acknowledgments

This research was performed in part at The Centre for Pathogen Genomics Innovation Hub, Department of Microbiology and Immunology, University of Melbourne at the Peter Doherty Institute for Infection and Immunity.

## Notes

https://github.com/rrwick/Are-reads-required

### Competing Interest Statement

The authors have declared no competing interest.

