## Supplementary figures for "Are reads required? High-precision variant calling from bacterial genome assemblies": Figure_S1.pdf

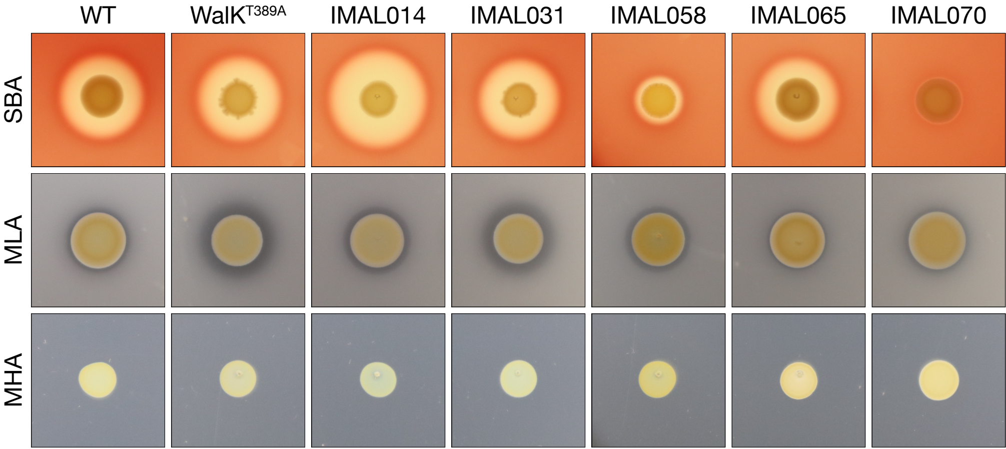


**Figure S1**: Phenotypic characterisation of the isolates.

SBA = Sheep blood agar; shows haemolytic activity.

MLA = *Micrococcus luteus* agar; shows the secretion/activity of one WalR-regulated peptidoglycan hydrolase (Atl).

MHA = Mueller-Hinton agar; shows the colour of the cells. IMAL058 is a SigB mutant, making it more yellow than the other isolates.
