## Supplementary figures for "Are reads required? High-precision variant calling from bacterial genome assemblies": Figure_S2.pdf

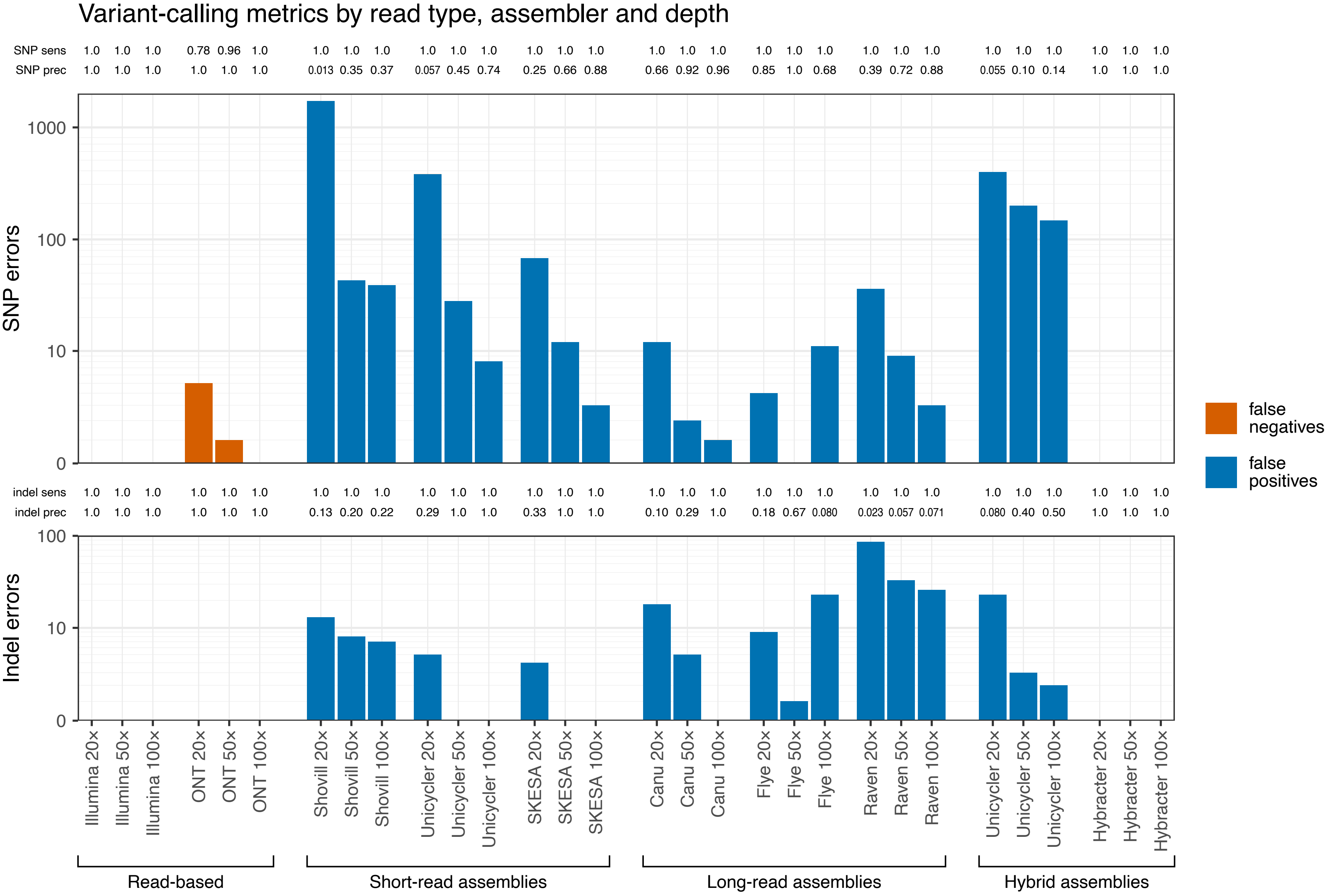


**Figure S2**: Variant calling metrics for both read- and assembly-based variant calling methods at each read depth. False negative and false positive errors are shown in the plots. Sensitivity (sens) and precision (prec) are shown above the plots. The y-axes have a pseudo-log transformation.
