## Supplementary figures for "Are reads required? High-precision variant calling from bacterial genome assemblies": Figure_S3.pdf

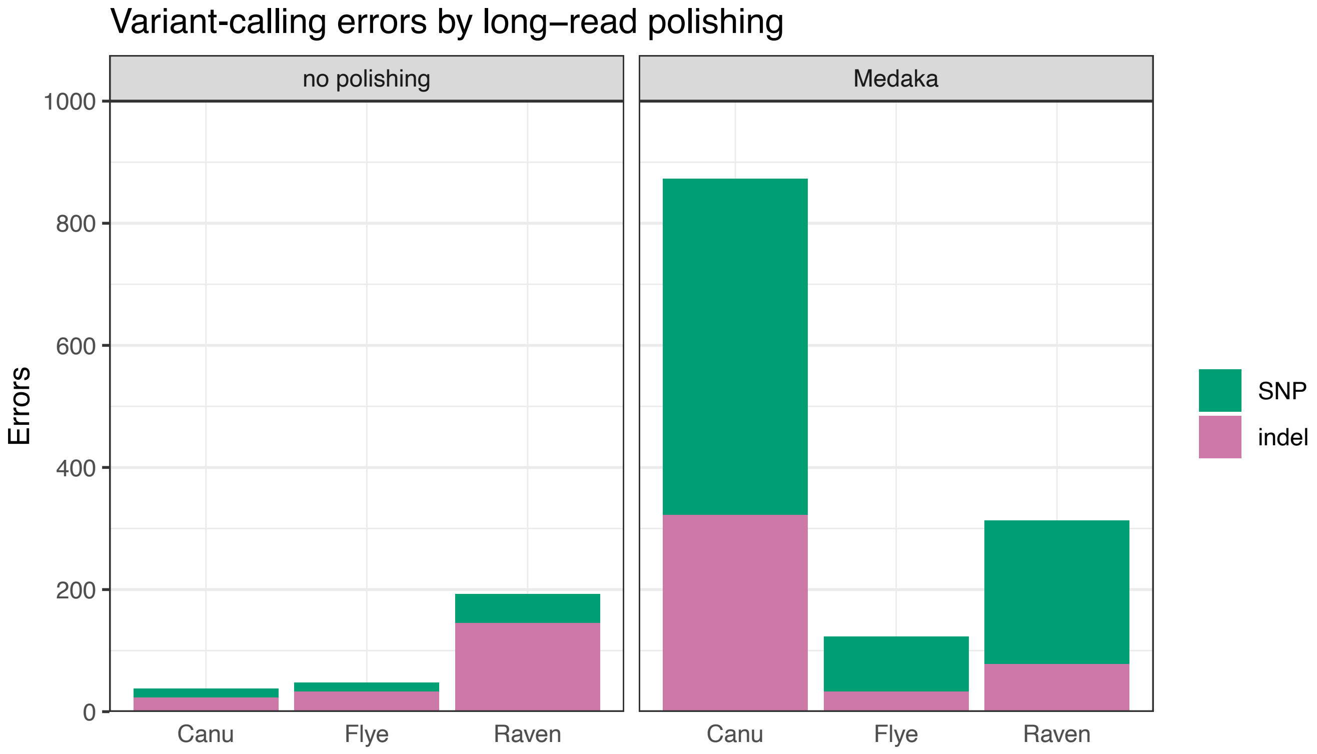


**Figure S3**: Variant-calling errors, before and after Medaka polishing, for each of the long-read assembly methods.
